# Missense mutations on SynGAP C2 domain impair membrane diffusion

**DOI:** 10.1101/2025.11.27.690967

**Authors:** Mattia Miotto, Leonardo Bo’, Giancarlo Ruocco, Silvia D’Angelantonio, Bernadette Basilico

## Abstract

*SYNGAP1* mutations have been linked to a range of neuropathological disorders and, more recently, to the insurgence of cancer. Despite its emerging relevance, the molecular basis of SynGAP’s action remains poorly understood. Here, using extensive molecular dynamics simulations integrated with structural analysis, we define how the SynGAP C2 domain associates with lipid bilayers. In particular, we observed spontaneous membrane binding of the domain in two distinct orientations, top and side, defined by the relative positioning of the C2 domain to its fold. These modes display markedly different dynamical properties: the top mode enables faster lateral diffusion on the membrane surface, whereas the side mode establishes more stable but less mobile contacts. Interestingly, pathogenic missense mutations mapping to the membrane-facing loops of SynGAP C2 disrupt these dynamics, leading to reduced diffusivity and altered membrane avidity. Our findings reveal a dual binding mechanism that underlies SynGAP’s membrane versatility and offer a general framework for how diverse mutations may perturb SynGAP activity across its polyedric biological functions.

## Introduction

SynGAP is a Ras-pathway GTPase-activating protein highly expressed at the postsynaptic density (PSD) of excitatory synapses [1, 2]. Here, SynGAP acts as a potent negative regulator of the RAS/ERK signaling and contributes to homeostatic control and plasticity [3, 4].

*De novo* heterozygous mutations in the *SYNGAP1* gene have emerged as a principal cause of the *SYNGAP1* syndrome, a neurodevelopmental disorder characterized by intellectual disability, epilepsy, autism spectrum disorder (ASD), and additional comorbidities [5, 6]. While the diagnosis of *SYNGAP1* -related disorder has expanded with advances in genomic sequencing, elucidating the molecular mechanisms underlying disease pathogenesis remains a major challenge.

The *SYNGAP1* gene encodes multiple protein isoforms that differ in their N- and C-terminal regions and display distinct expression patterns during development and across brain regions [7, 8], suggesting a multifaceted role that extends beyond synaptic signaling. This diversity complicates the functional interpretation of genetic variants and the design of effective model systems. Nevertheless, all isoforms share a conserved core structure comprising a C2 domain and the RasGAP domain, implicating these regions as central to SynGAP’s function. The C2 domain is broadly conserved among several proteins involved in signal transduction and membrane trafficking, distinguished by its adaptable lipid-binding properties, including interactions with key membrane components such as phosphatidylserine and phosphatidylcholine [9]. The coordinated action of the C2 and RasGAP domains is required for SynGAP’s GTPase-activating function [10]. *In vivo* studies using rat models with targeted deletions in the SynGAP C2 and GAP domains demonstrated their importance in the appearance of key pathological aspects previously observed in *SYNGAP1* haploinsufficiency models, such as cognitive deficits, altered social behaviors, and increased seizure susceptibility [11]. These observations underscore the functional importance of the C2 domain in maintaining synaptic integrity and neurological function.

Despite these advances, a comprehensive understanding of how specific missense mutations within the C2 domain influence SynGAP membrane-binding properties and downstream signaling remains under-investigated. Given the observed clinical heterogeneity among individuals with non-truncating *SYNGAP1* mutations, there is an urgent need for robust mechanistic frameworks to interpret the pathogenicity of these variants. Previous atomistic modeling of SynGAP’s N-terminal region (PH–C2–GAP) established an upright (“loop legs”) membrane binding mode by aligning the C2 domain onto membrane-bound templates from structurally analogous proteins. This approach characterized the stability assuming a given orientation rather than observing spontaneous membrane association or alternative binding modes in unbiased simulations [13]. Furthermore, the effects of disease-associated missense mutations on C2 domain dynamics at the membrane remained largely inferred rather than directly quantified. Here, we address this gap by integrating extensive molecular dynamics simulations with targeted mutagenesis to systematically evaluate patient-specific missense mutations identified in the C2 domain. Our approach enables detailed analysis of whether and how these variants alter C2 domain structure and membrane interaction, providing new insights into the role of non-truncating *SYNGAP1* variants in neurodevelopmental disorders. Finally, our findings highlight the frequently overlooked pathogenic potential of missense mutations and offer a foundation for improved functional annotation and precision diagnostics in *SYNGAP1* -related disorders.

## Results

### SynGAP C2 domain adopts a type II fold

At present, no experimentally solved atomic structure of human SynGAP protein is available. As starting point for our computational investigation, we considered the structure predicted by AlphaFold [12]. The structure shows three ordered regions corresponding to the PH, C2 and GAP domains followed by an extended disordered region. Notably, the C2 domain assumes a type II fold, characterized by the orientation of the C- and N-termini with respect to the rest of the domain and a specific organization of the *β*-sheets (see Figure 1a). As our interest lies in the stability and function of the C2 domain, we initially neglected the disordered region, truncating the structure at residue 740. Thus, we assumed that the stability of the domain is not affected by the presence of the disordered region. Next, we probed the stability of the SynGAP ordered region via full-atom molecular dynamics (MD) simulations. In particular, we ran two MD simulations considering SynGAP residue region 150-740 (referred as full ordered region, ‘full’) and region 253-413 (referred as C2 domain, ‘C2’). We first evaluated the behavior of the Root Mean Square Displacement (RMSD) and Gyration radius (Gy) as a function of the simulation time for the C2 domain and we compared its equilibrium distribution values with the full protein (Figure 1b-c). Both descriptors vary in the same ranges indicating that the predicted domain is stable in the simulated time scale. To evaluate whether the isolated C2 domain reproduces the local structural dynamics it displays within the full SynGAP protein, we compared its conformational fluctuations under the two simulation conditions. Analysis of the residue-wise RMSF profiles revealed highly similar patterns of flexibility between the C2 domain simulated alone in water and the same region embedded in the complete SynGAP structure (Fig. 1d). The correspondence in fluctuation amplitudes across key loops and secondary-structure elements indicates that the intrinsic dynamical features of the C2 domain are largely maintained even outside the context of the full protein. Notably, the glycine-rich loop (residues 370-395) exhibits the highest fluctuations (see also Figure 1e).

**FIG. 1:**
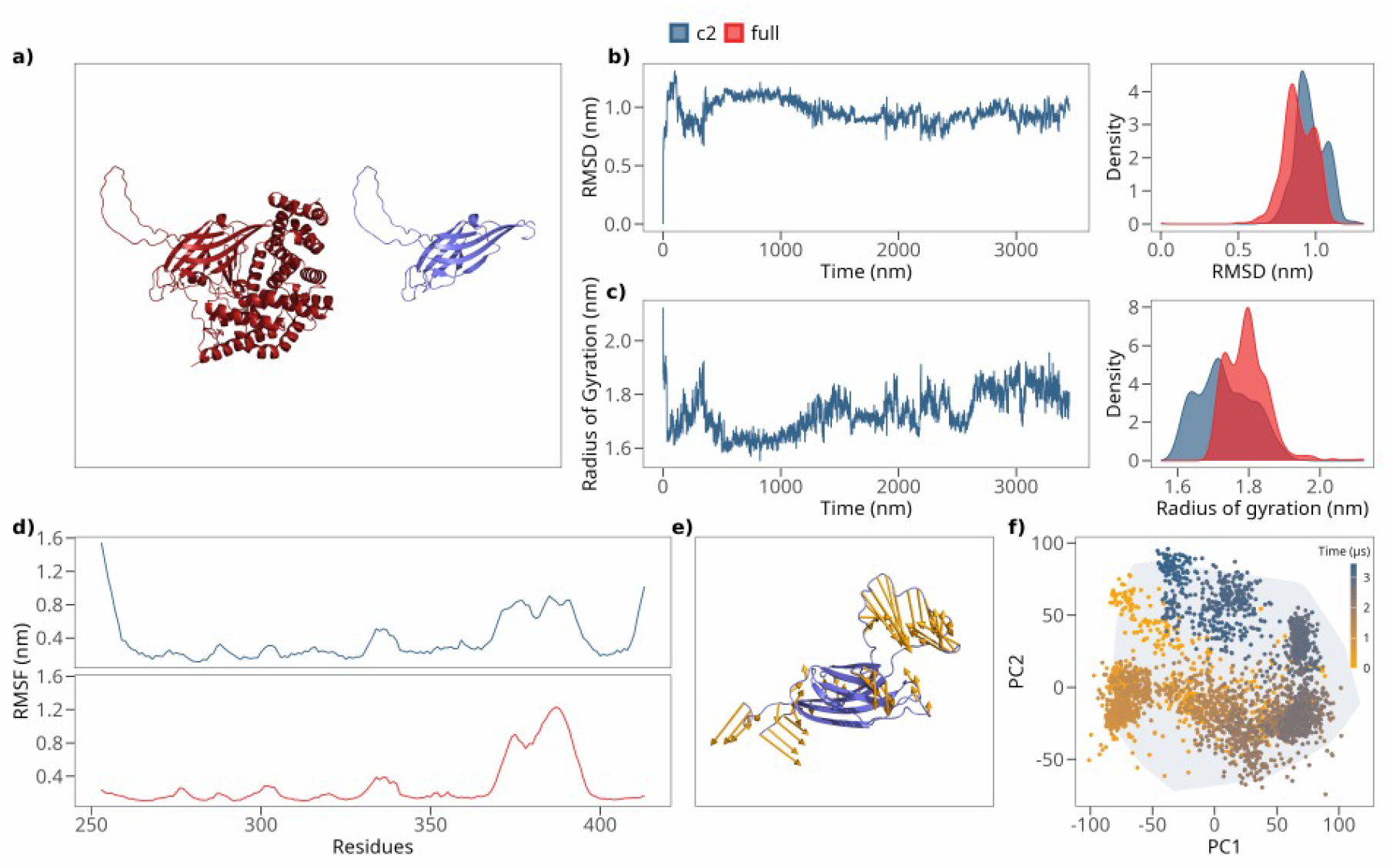
Dynamical features of the SynGAP C2 domain. **a)** Cartoon representation of the 3D structure of SynGAP as predicted by AlphaFold [12]. The C2 domain is predicted to assume a type II fold. **b)** Root mean square displacement (RMSD) as a function of the simulation time for the SynGAP C2 domain in water (left) and distributions of the RMSD in the equilibrium regime for the C2 domain alone and embedded in the full SynGAP protein (right). **c)** Same as in panel b) but for the radius of gyration. **d)** Root mean square fluctuations (RMSF) as a function of the C2 domain residues. Top panel displays the RMSF for the C2 domain alone in water, while bottom panel shows the RMSF values for the C2 domain embedded in the whole SynGAP structure. **e)** Graphical representation of the motion of the C2 domain in terms of its normal modes. Yellows arrows represents the direction and intensity of the fluctuations. **f)** Projection of the trajectory of the molecular dynamics simulation of the C2 domain in the plane identified by the two principal components of the configurations covariance matrix. Colored dots represent the position of the configuration when projected in the plane. Colors ranges from yellow to blue as simulation time increases. The region explored by the C2 domain embedded in the full SynGAP protein is marked in grey.

To further compare the conformational landscapes sampled under the two conditions, we projected the molecular dynamics trajectory of the isolated C2 domain onto the first two principal components of its covariance matrix (see Methods). The distribution of sampled configurations (Fig. 1f) overlaps extensively with the region explored by the C2 domain when part of the full SynGAP protein, delineated in grey. This overlap indicates that the isolated domain traverses a conformational ensemble that is highly consistent with that of the C2 region in its native protein environment. Together, these analyses demonstrate that the isolated C2 domain captures the essential dynamical behavior of the C2 region within SynGAP, supporting the use of the isolated construct as a reliable model for probing C2-specific structural and functional properties.

### SynGAP C2 domain is predicted to bind lipid bilayers via two distinct modes

To characterize the membrane-interacting behavior of the SynGAP C2 domain, we performed a set of 2*µs*-long molecular dynamics simulations of a system comprising the C2 domain and a lipid bilayer, that mimicks the cellular membrane. The C2 domain was placed in a randomly selected orientation with respect to the membrane plane and at a distance higher than 12 *A*, so that no direct interactions are present between the domain and the membrane at the initial step of the dynamics. The composition of the membrane has been set along the lines of Larsen *et al*. [14] (see Figure 2a and Methods for more details). After a transient time of about 20 *ns*, SynGAP C2 domain spontaneously binds to the lipid membrane, adopting two recurrent and stable binding configurations. Indeed, across 20 independent replicas, 5 the protein domain was consistently observed to associate with the membrane surface, either through its lateral surface (‘Side’ mode) or through the top face of the *β*-sandwich (‘Top’ mode). Figure 2b displays the time evolution of the distance between the protein center of mass. In the ‘Top’ mode (red trace), the C2 domain approached the lipid bilayer perpendicularly and stabilized with its principal axis roughly normal to the membrane plane, as reflected in the orientation angle plot (Fig. 2b, middle). In contrast, the ‘Side’ mode (blue trace) displayed a tilted orientation with the long axis parallel to the lipid bilayer, remaining stably adsorbed throughout the simulation. Residue-wise minimum distance analysis (Fig. 2b, bottom) revealed that distinct regions of the C2 domain mediate the two interactions. In the ‘Top’ mode, the membrane-proximal residues were predominantly localized on loops at the tip of the *β*-sandwich, while in the ‘Side’ mode, a broader surface involving *β*-strands and adjacent loops maintained contact with the lipid headgroups. Clustering of mean minimal distances across all replicas (Fig. 1c) confirmed that these two contact patterns represent the dominant binding configurations sampled during the simulations, with the ‘Side’ configuration being the more probable.

**FIG. 2:**
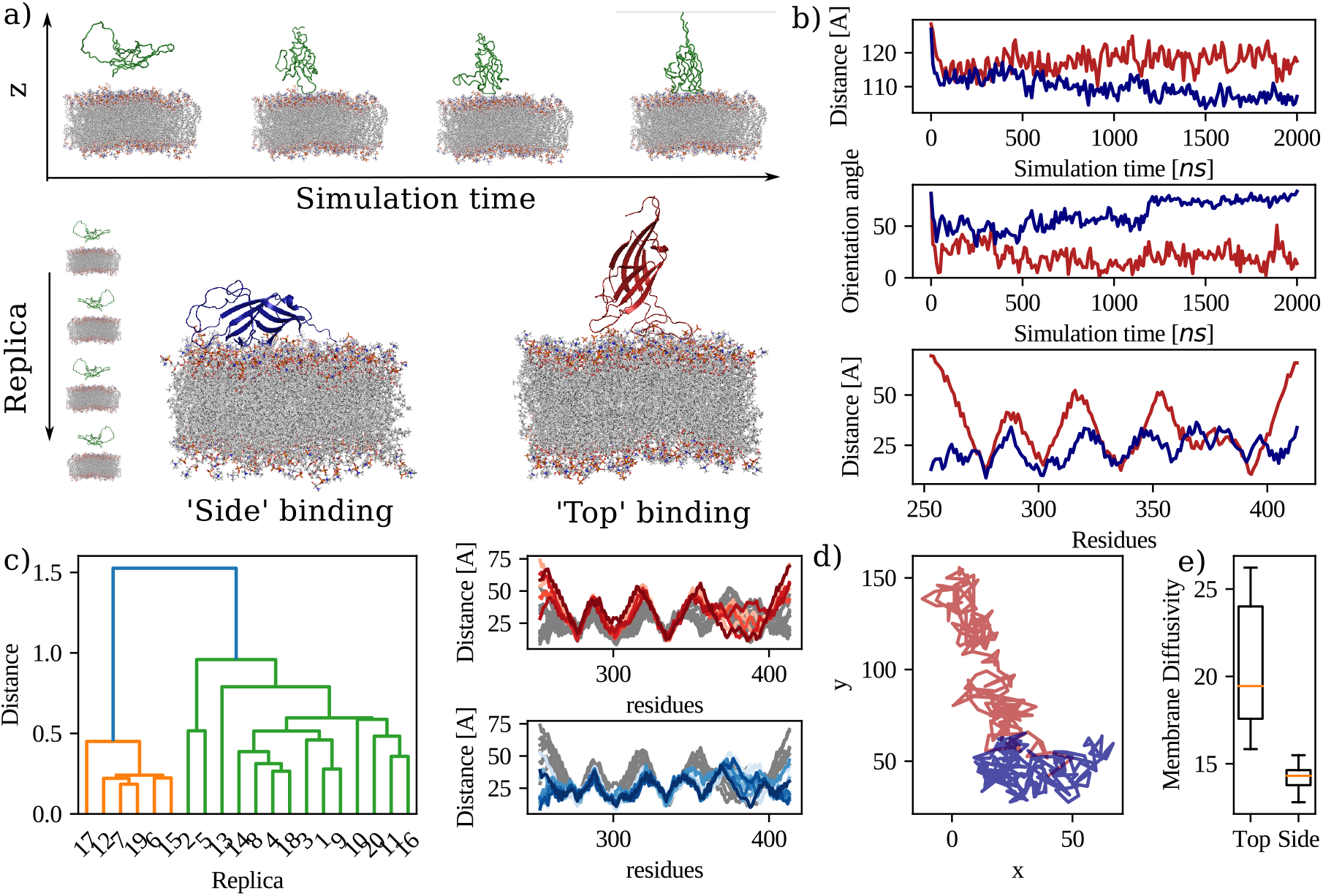
SynGAP C2 domain-membrane binding modes. **a)** Snapshot of the time evolution of the molecular dynamics simulation for a system composed of the SynGAP C2 domain and a lipidic membrane. The C2 domain spontaneously binds to the membrane in two different modes. **b)** From top to bottom, distance of the protein center of mass from the center of the lipidic bilayer as a function of the simulation time for two representative runs where the C2 domain binds in ‘Side’ (blue) and ‘Top’ (red) configuration; Orientation angle of the principal axis of the C2 domain as a function of the simulation time; Mean minimal distance after binding of each residue of the C2 domain from the membrane. **c)** Clustering analysis of the mean minimum distances for all 20 replica of the binding process (orange = Top mode; green = Side mode). **d)** Projection of th trajectory of the C2 center of mass in the membrane plane for two representative MD simulations. **e)** Box plot representation of log-transformed diffusion coefficients for the two binding modes.

### Binding modes show different dynamical properties

Analysis of the two-dimensional trajectories of the C2 domain center of mass in the membrane plane indicated distinct diffusive behaviors for each mode (Fig. 2d). The “Top” configuration exhibited more confined lateral motion, consistent with deeper insertion or stronger electrostatic anchoring, whereas the “Side” configuration showed broader lateral diffusion. Quantification of the diffusion coefficients (Fig. 2e) revealed a significantly lower median diffusion for the “Top” mode, as evidenced by the box plot distribution of log-transformed diffusion values.

Together, these results demonstrate that the SynGAP C2 domain binds to membranes via two distinct, stable modes that differ in orientation, contact residues, and lateral mobility, suggesting that each configuration may play a specific functional role in membrane association dynamics.

### Missense mutations on C2 domain impair membrane interactions

To investigate how disease-associated missense mutations affect the structural and dynamical properties of the SynGAP C2 domain, we compared the dynamical behavior observed in extensive molecular dynamics simulations for the wild-type (WT) C2 domain and a set of pathogenic/likely pathogenic missense *SYNGAP1* variants distributed across the C2 (Fig. 3a).

**FIG. 3:**
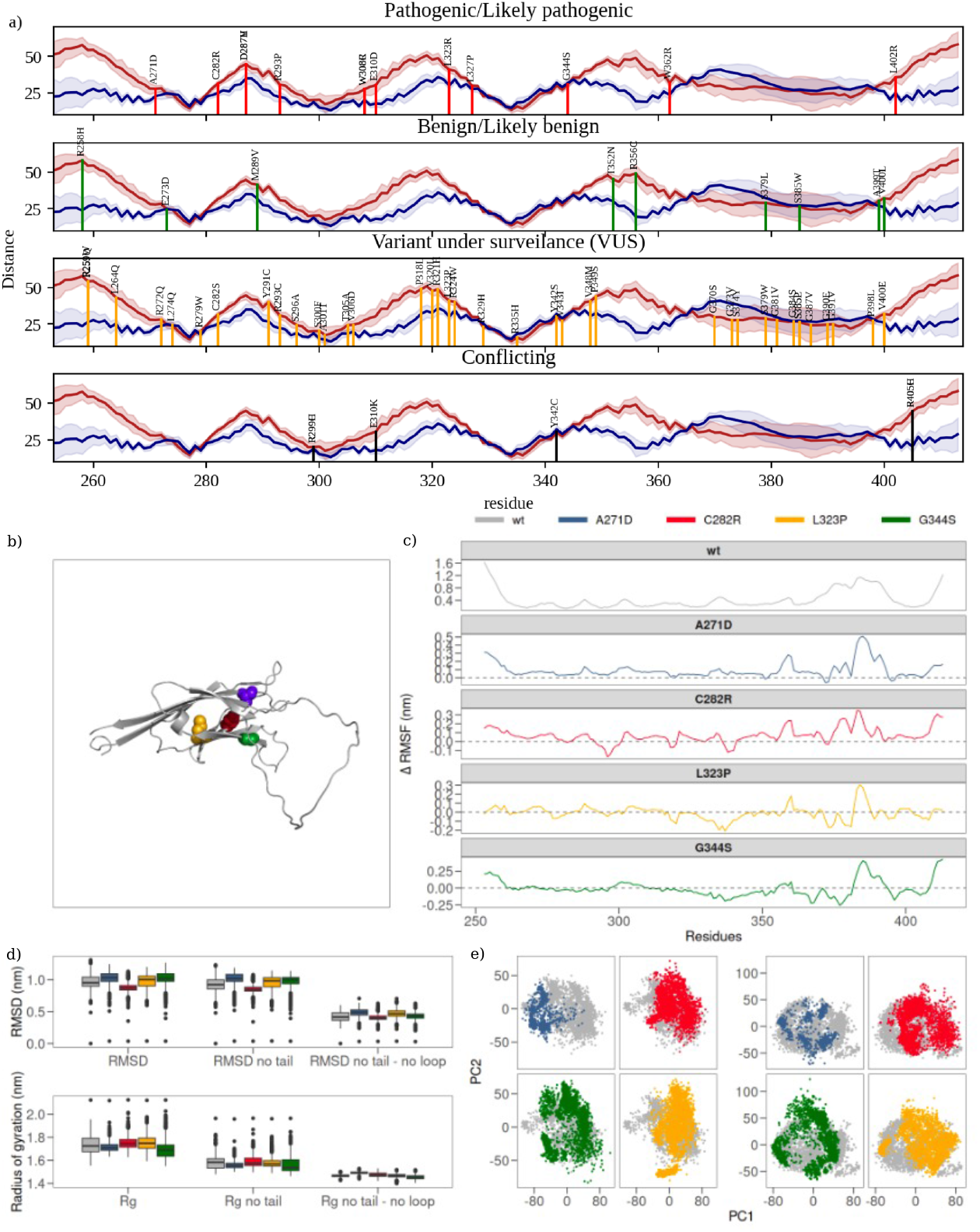
Dynamical features of the considered non-truncating *SYNGAP1* variants. **a)** Localization of the known missense mutations on the SynGAP C2 domain stratified by clinical severity. **b)** Cartoon representation of the C2 domain with the considered mutated residues highlighted in different colors. **c)** From top to bottom, root mean square fluctuations (RMSF) as a function of the WT C2 domain residues and difference between the RMSF of the considered variants with respect to the WT. **d)** Box plots of RMSD and Gyration ratios distributions at the equilibrium regime for the C2 domain in the WT and the considered variants (see legend in panel c). From left to right, whole C2 domain, domain excluding the N- and C-terminal residues (residues 253-260 and 405-413; and domain excluding the N- and C-terminal residues and larger loop (residues 370-395). Left panels, projections of MD trajectories for C2 domain missense variants onto the plane defined by the first two principal components (PC1-PC2) of the configuration covariance matrix. Colored dots represent the position of the configuration when projected in the plane. The region explored by the WT C2 domain is marked in grey. Right panels consider only the larger loop residues for the covariance matrix calculation.

To do so, we used the public online database ClinVar [15] to collect known missense variants mapping to the C2 for a total of 60 variants stratified by reported clinical severity. First, we examined how the mutated residues are distributed among the C2 domain’s structural elements and mapped these positions onto the C2–membrane distance profiles (Figure 3a). Examining the positional distribution of variants and their clinical severity, we found no clear correlation, indicating no pathogenicity hotspot within the C2 domain. Notably, no known pathogenic/likely pathogenic mutations are observed on the residues we predict to directly mediate the C2 binding with the membrane. On the contrary, among mutations that are currently under surveilance (VUS), mutations A301T and R335H involve residues found in interaction with the lipid membrane in the WT simulations. To understand the effect of mutations on the C2 domain, we selected a set of representative variants reported in Table I, considering severity and proximity to the membrane binding regions.

**TABLE I:** Missense mutations identified in the C2 domain and studied in this work.

| SYNGAP1 mutation | Severity | Top Diffusivity | SYNGAP1 mutation | Severity | Side Diffusivity |
| --- | --- | --- | --- | --- | --- |
| WT | - | $22 \pm 1$ | WT | - | $13.4 \pm 0.5$ |
| A271D | pathogenic | $17 \pm 2$ | C282R | pathogenic | $13.1 \pm 0.7$ |
| C282R | pathogenic | $19 \pm 4$ | L323P | pathogenic | $14.8 \pm 0.4$ |
| A301T | VUS | $14 \pm 1$ | R335H | VUS | $11.6 \pm 0.6$ |
| G344S | pathogenic | $20 \pm 1$ | G344S | pathogenic | $12.2 \pm 0.6$ |

Residue-level flexibility analysis revealed that all variants preserved the overall C2 domain fold but exhibited local alterations in mobility (Fig.3c). Specifically, the root mean square fluctuation (RMSF) profiles showed that mutations near loop regions or at membrane-interacting surfaces increased local flexibility relative to WT. The different RMSF plots highlighted mutation-specific effects: variants affecting residues at the *β*-sandwich edges or in the loop connecting *β*-strands showed enhanced fluctuations, while mutations in *β*-sandwich core displayed dampened motions, suggesting local rigidification (see e.g. L323P and G344S).

Quantitative analysis of structural stability through root mean square deviation (RMSD) measurements in the equilibrium regime (Fig.3d) confirmed that the global C2 domain structure remained stable across all variants, with median RMSD values comparable to the WT. However, when the flexible N- and C-terminal segments were excluded, the spread of RMSD distributions decreased markedly, underscoring that most of the structural variability arises from peripheral residues. Further exclusion of the larger loop region resulted in nearly identical RMSD distributions among all systems, indicating that the core *β*-sandwich scaffold remains unaffected by point mutations.

Consistent trends were observed for the radius of gyration (Fig.3d). The variants maintained compactness comparable to the WT, with only minor differences in the distributions once terminal and loop regions were removed. These results suggest that the overall folding stability of the C2 domain is largely preserved, while the local dynamics around flexible regions are most sensitive to mutation.

Principal component analysis (PCA) of the conformational ensembles provided further insight into collective motions (Fig. 3e). Projection of the trajectories onto the first two principal components revealed that the WT explored a well-defined region of conformational space (grey area), while most variants sampled broader or shifted regions, indicating altered conformational preferences. When the analysis was restricted to the large loop residues, these differences became more pronounced, with some variants sampling configurations outside the WT’s conformational basin, consistent with enhanced loop flexibility or alternative conformations.

Finally, to assess the functional implications of these dynamical alterations, we evaluated how different *SYNGAP1* missense variants affect the diffusivity of the C2 domain in the two membrane-bound configurations. Diffusivity for all considered variants are reported in Table I. While mutations tend to reduce diffusivity, A301T and R335H exhibited markerly reduced diffusivity relative to the WT, suggesting stronger or more persistent membrane contacts.

Overall, these findings indicate that non-truncating *SYNGAP1* mutations modulate the local flexibility and conformational dynamics of the C2 domain without disrupting its global architecture. The resulting alterations in loop motion and membrane diffusivity may underlie variant-specific effects on SynGAP membrane association and signaling behavior (see Figure 4 for a graphical representation).

**FIG. 4:**
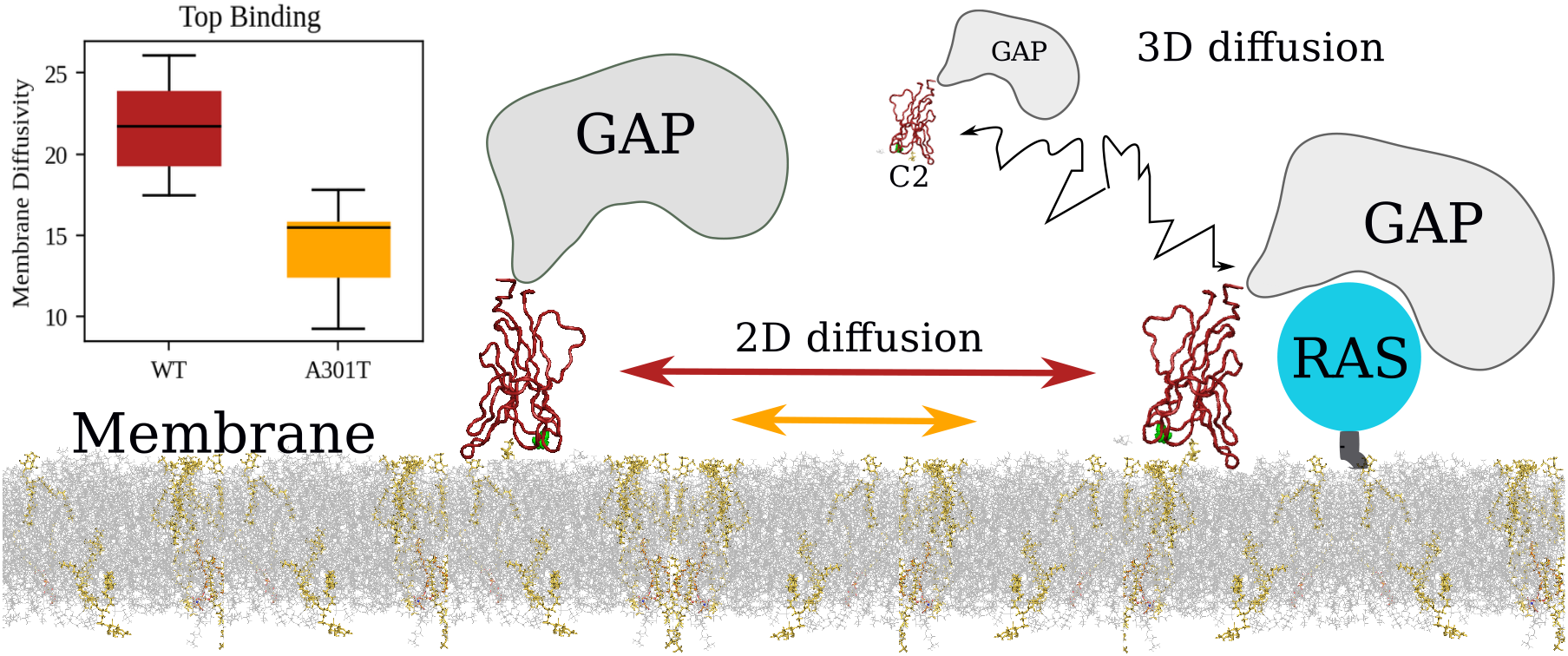
Effect for missense mutations on the membrane binding sides. Sketch of the suggested role of the C2 domain in mediating SynGAP membrane binding and subsequent 2D diffusion toward RAS proteins. In the inset, box plots of the diffusivity distribution for the WT and A301T variant when found in the ‘Top’ binding mode.

### Differential lipid-residue interactions accounts for the observed diffusivities

Finally, we analysed the specific interactions between the C2 domain and the lipid bilayer to assess why pathogenic variants display differences in their interaction with the membrane. In particular, we evaluated the lipid–protein contact probabilities stratified by the main three classes of lipids present in the membrane: POPC, POPI and POPS. Comparison of the wild-type (WT) and *SYNGAP1* A301T variant revealed a clear increase in membrane engagement for the mutant. Across the entire C2 sequence, A301T displayed consistently higher contact probabilities with all three lipid species compared to the WT domain (Fig. 5b). This global enhancement in lipid contacts suggests that the A301T substitution promotes a generally more adhesive or stable association with the membrane surface. Indeed, visual inspection of the C2-lipid disposition, shows that the loop containing residue 301 direcly interact with POPC lipids, while the same loop in the WT is not found in proximity of those lipids (see snapshots in Figure 5).

**FIG. 5:**
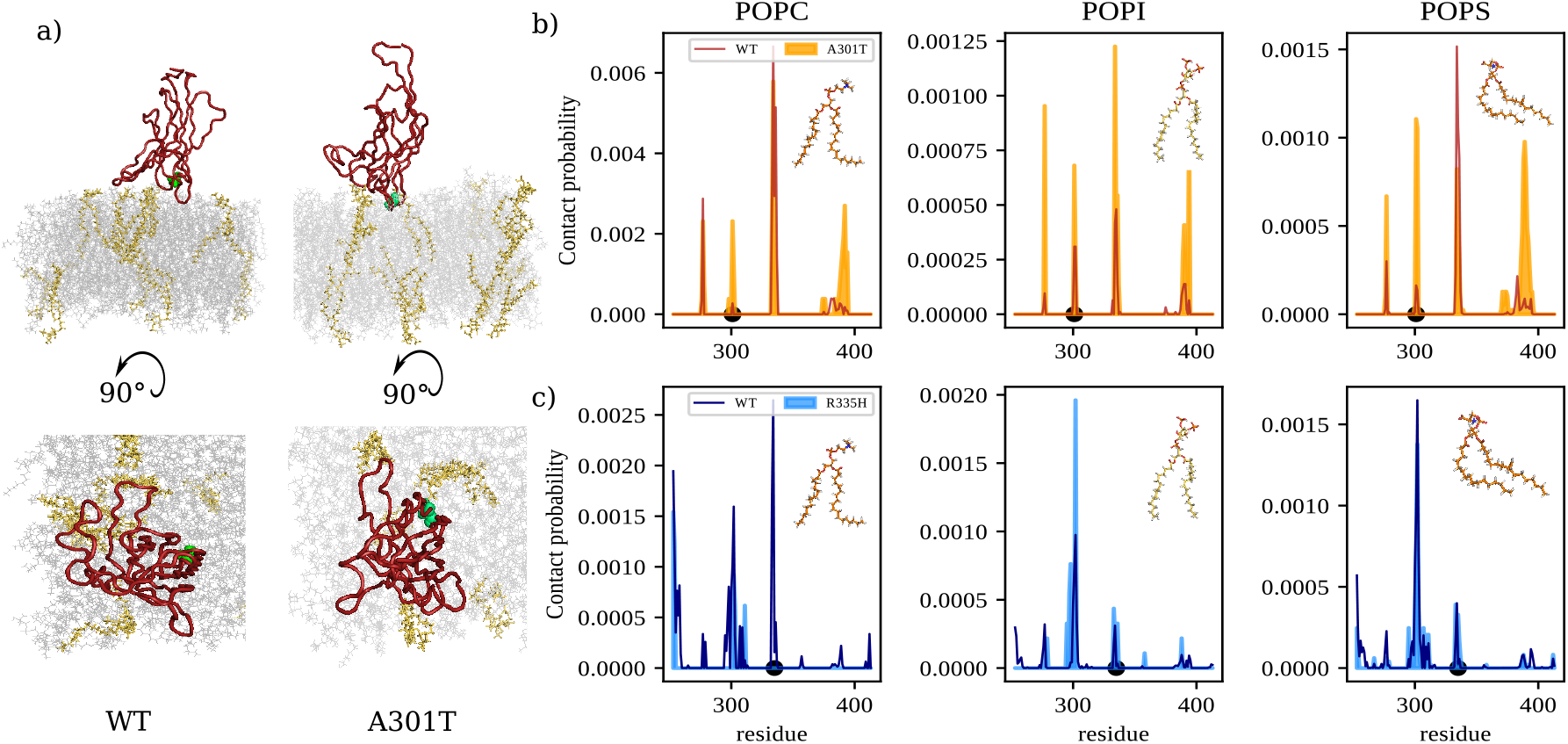
Differential interaction between C2 and membrane lipids. **a)** Snapshot from the time evolution of the molecular dynamics simulation for a system composed of the SynGAP C2 domain and a lipidic membrane, with the C2 domain bound to the membrane. POPI lipids are highlighted in yellow, while residue 301 is marked in green. **b)** Contact probabilities of the WT (red) and variant A301T (orange) C2 domain as a function of the C2 residues toward the three kinds of lipids present in the simulated membrane. **d)** Same as for panel b) but considering the WT and variant R335H when found in the ‘Side’ binding mode.

In contrast, the R335H variant showed a distinct and lipid-specific alteration in its interaction pattern when the domain adopted the “Side” binding mode (Fig. 5c). Relative to WT, R335H exhibited a reduction in contacts with POPC lipids, indicating weaker engagement with the bulk, zwitterionic component of the bilayer. However, this decrease was accompanied by a notable increase in contacts with POPI, pointing to a shift in lipid preference toward the anionic PI species. This redistribution suggests that the R335H substitution modifies the electrostatic landscape of the membrane-binding interface, biasing interactions toward negatively charged lipids while diminishing those with more neutral membrane components.

Together, these analyses highlight how different mutations in the C2 domain can modulate membrane affinity either globally (A301T) or in a lipid-selective manner (R335H), potentially altering the stability and specificity of SynGAP membrane association.

## Discussion

In this study, we used extensive molecular dynamics simulations to characterize membrane binding by the SynGAP C2 domain and to evaluate the potential disruptive effects of different *SYNGAP1* missense variants. We found that the C2 domain binds the membrane in two distinct orientations that differ markedly in their lateral mobility on lipid bilayers, as well as in their sensitivity to the location of variants within the membrane-interacting loops. The ‘top’ binding mode exhibits faster lateral diffusion, consistent with a more transient and dynamic interaction with the membrane, whereas the ‘side’ mode forms more stable but less mobile contacts. Mutations in the membrane-facing loops substantially reduce diffusivity, indicating that these regions are crucial for maintaining the flexible association required for rapid membrane exploration.

Notably, our findings align with prior structural and biophysical studies showing that C2 domains can adopt diverse lipid-binding geometries and dynamics depending on domain topology and membrane lipid composition. In particular, multi-scale molecular dynamics studies of *Ca*^+2^-independent C2 domains have reported distinct ‘top’ and ‘side’ binding modes governed by the distribution of charged residues and interactions with anionic lipids such as phosphatidylserine and phosphoinositides [14]. Our observation of these two orientations for SynGAP’s C2 domain is therefore in line with these general principles. The distinct lateral mobilities of the two modes are particularly intriguing. Experimental single-molecule studies of membrane-targeting domains have reported diffusive regimes ranging from Brownian to super-diffusive motion, depending on the depth and duration of membrane engagement [16, 17]. The faster-diffusing top mode identified here may correspond to a shallow search configuration that enables efficient sampling of the membrane surface, whereas the more stable side mode could reflect a transition to a functionally engaged state, possibly preceding effector recruitment or catalysis.

Our data also provide mechanistic insight into how the C2 domain regulates SynGAP’s GTPase-activating function. The C2 domain is known to be essential for the Ras/RapGAP activity of SynGAP: constructs containing both the C2 and GAP domains display full catalytic efficiency, whereas the GAP domain alone is inactive [10]. Beyond accelerating GAP activity, the C2 domain likely contacts the Ras allosteric lobe [18]. It is plausible that the membrane-binding geometry modulates this activity by influencing orientation and proximity to the membrane, thereby controlling substrate access and catalytic turnover. We hypothesize that, by localizing SynGAP to membranes, the C2 domain enables lateral diffusion and functional exploration of signaling partners (such as Ras) within defined membrane microdomains. In synapses, it may help orchestrate synaptic composition and plasticity together with PSD-95 scaffolding and post-translational regulation.

Altogether, our findings propose a dual binding mechanism through which SynGAP may achieve both dynamic membrane scanning and stable engagement, depending on functional context. Perturbations that shift this balance, such as pathogenic or cancer-associated variants, may underlie diverse disease mechanisms. Future work should test these predictions experimentally by examining how membrane composition, post-translational modifications, or disease variants influence the equilibrium between top and side binding, and by assessing whether changes in membrane diffusivity translate into altered Ras/Rap or Wnt/*β*-catenin signaling outputs.

## Materials and Methods

### Dataset

Three-dimensional structure of the SynGAP protein has been retrieved by AlphaFold predicted human proteome [12]. The C2 domain has been extracted considering residues from 253 to 413. *SYNGAP1* variants (see Table I) have been obtained via computational mutagenesis using PyMol suite on the WT predicted structure.

### Molecular dynamics simulations

All simulations were performed using Gromacs [19]. Topologies of the system were built using the CHARMM-36 force field [20]. For simulations without lipidic by-layers, the protein was placed in a dodecahedric simulative box, with periodic boundary conditions, filled with TIP3P water molecules [21]. For all simulated systems, we checked that each atom of the proteins was at least at a distance of 1.1 nm from the box borders. Each system was then minimized with the steepest descent algorithm. Next, a relaxation of water molecules and thermalization of the system was run in NVT and NPT environments each for 0.1 ns at 2 fs time-step. The temperature was kept constant at 300 K with v-rescale thermostat [22]; the final pressure was fixed at 1 bar with the Parrinello-Rahman barostat [23].

LINCS algorithm [24] was used to constraint bonds involving hydrogen atoms. A cut-off of 12 Å was imposed for the evaluation of short-range non-bonded interactions and the Particle Mesh Ewald method [25] for the long-range electrostatic interactions. The described procedure was used for all the performed simulations.

#### A. Protein-membrane simulations

We used CHARMM-GUI to construct a membrane system composed of a lipid bilayer and the C2 domain. The bilayer lies in the xy plane, and the C2 domain was translated by 40 Å along the z axis with respect to the bilayer midplane.

A parallelepiped simulation box was employed, imposing a minimum distance of 22.5 Å between any atom in the system and the nearest box face. CHARMM-GUI determined the box and the membrane dimensions. Each leaflet of the bilayer was composed of 90 POPC molecules, 17 POPS molecules, 3 POPI(2,5)P molecules, and 3 POPI(2,4)P molecules [14].

The simulation box was neutralized adding *Na*+ ions. The system was minimized (steepest-descent algorithm), equilibrated in the NVT ensemble for 125 ps using the V-rescale thermostat, and subsequently equilibrated in the NPT ensemble for 125 ps using the C-rescale barostat with semi-isotropic pressure coupling. LINCS constraints were applied throughout, together with additional positional restraints during equilibration. The production simulations were run at T = 300 K, with a time step of 1 fs during minimization/equilibration and 2 fs during production.

### Statistics and Reproducibility

All molecular dynamics simulations of the C2 domain (WT and mutated) were 2000 ns long, which guarantees that every system has reached equilibrium conformations, and performed in three replica. C2-membrane simulations were performed for a time of 1000 ns, each repeated for 5 replicas. WT has repeated for 20 replicas to achieved statistical significancy for the binding poses determination.

### Principal component analysis and clustering

We performed a principal component analysis (PCA) in which the starting matrix consisted of rows equal to the number of residue and columns equals to the sampled configurations. The clustering analysis was performed using the ‘dendogram’ function of Python, preserving the default clustering algorithm.

#### 1. Diffusion coefficient

The diffusion coefficient *D* quantifies long-time translational motion and is computed from the mean squared displacement (MSD),

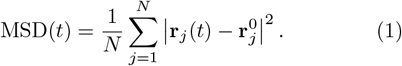

The diffusion coefficient is defined as

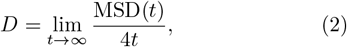

in two spatial dimensions [26].

## Data Availability

The data that support the findings of this study are available from the corresponding author upon reasonable request.

## Acknowledgements

We thank the APS Famiglie Syngap1 Italia and APS Le Note di Fra for supporting the *SYNGAP1* research (B.B.). Additionally, G.R. received fundings by grants from ERC-2019-Synergy Grant (ASTRA, n. 855923); EIC-2022-PathfinderOpen (ivBM-4PAP, n. 101098989); Project ‘National Center for Gene Therapy and Drugs based on RNA Technology’ (CN00000041) financed by NextGeneration EU PNRR MUR—M4C2—Action 1.4—Call ‘Potenziamento strutture di ricerca e creazione di campioni nazionali di R&S’ (CUP J33C22001130001); Project ECS 0000024 Rome Technopole-CUP B83C22002820006, PNRR Missione 4 Componente 2 Investimento 1.5, finanziato dall’Unione europea – NextGenerationEU; Seed Sapienza (SP1221842D4094CE) (B.B. and S.D.A.).

